# scMEGA: Single-cell Multiomic Enhancer-based Gene Regulatory Network Inference

**DOI:** 10.1101/2022.08.10.503335

**Authors:** Zhijian Li, James S Nagai, Christoph Kuppe, Rafael Kramann, Ivan G Costa

## Abstract

The increasing availability of single-cell multi-omics data allows to quantitatively characterize gene regulation. We here describe scMEGA (Single-cell Multiomic Enhancer-based Gene Regulatory Network Inference) to infer gene regulatory networks by combining single-cell gene expression and chromatin accessibility profiles. This enables to study of complex gene regulation mechanisms for dynamic biological processes, such as cellular differentiation and disease-driven cellular remodeling. We provide a case study on gene regulatory networks controlling myofibroblast activation in human myocardial infarction

## Background

Single-cell RNA sequencing (scRNA-seq) and ATAC-seq (scATAC-seq) techniques provide an unprecedented opportunity to understand gene regulation at the single-cell level by capturing orthogonal molecular information [1], i.e., gene expression by scRNA-seq and activity of regulatory elements and transcription factors (TFs) by scATAC-seq [2].Applying both assays on the same biological samples generates single-cell multi-omics data which allow to computationally infer cell-type-specific gene regulatory networks (GRN) for different cellular systems, such as fly brain development [3], human myocardial infarction [4], and human brain development [5]. However, such analysis is usually based on complex bioinformatic pipelines that require different tools for each of the steps, such as Seurat for scRNA-seq analysis and data integration [6], ArchR for scATAC-seq analysis and trajectory inference [7], chromVAR for TF activity estimation [8] and igraph [9] for general network analysis.

Currently, two computational tools, Pando [5] and CellOracle [10], are available for GRN inference based on single-cell multi-omics profiles. Pando, however, focus on the identification of regulatory networks limited to TF-TF interaction and does not provide methods for modality integration or trajectory analysis. CellOracle needs to first assemble a basic GRN structure based on reference scATAC-seq data and requires several pre-processing steps for both scRNA-seq and scATAC-seq analysis, which limits its usage.

We therefore developed scMEGA as a general framework to quantitatively infer GRN by taking single-cell multi-omics profiles as input. scMEGA enables an end-to-end analysis of multi-omics data for GRN inference including modalities integration, trajectory analysis, enhancer-to-promoter association and network analysis and visualization (Fig.1). For this, it provides new functionalities and combines some existing methods from Seurat, ArchR, chromVAR and igraph. It is implemented as an R package and is working on Seurat objects, making it compatible with the Seurat ecosystem.

**Figure 1.**
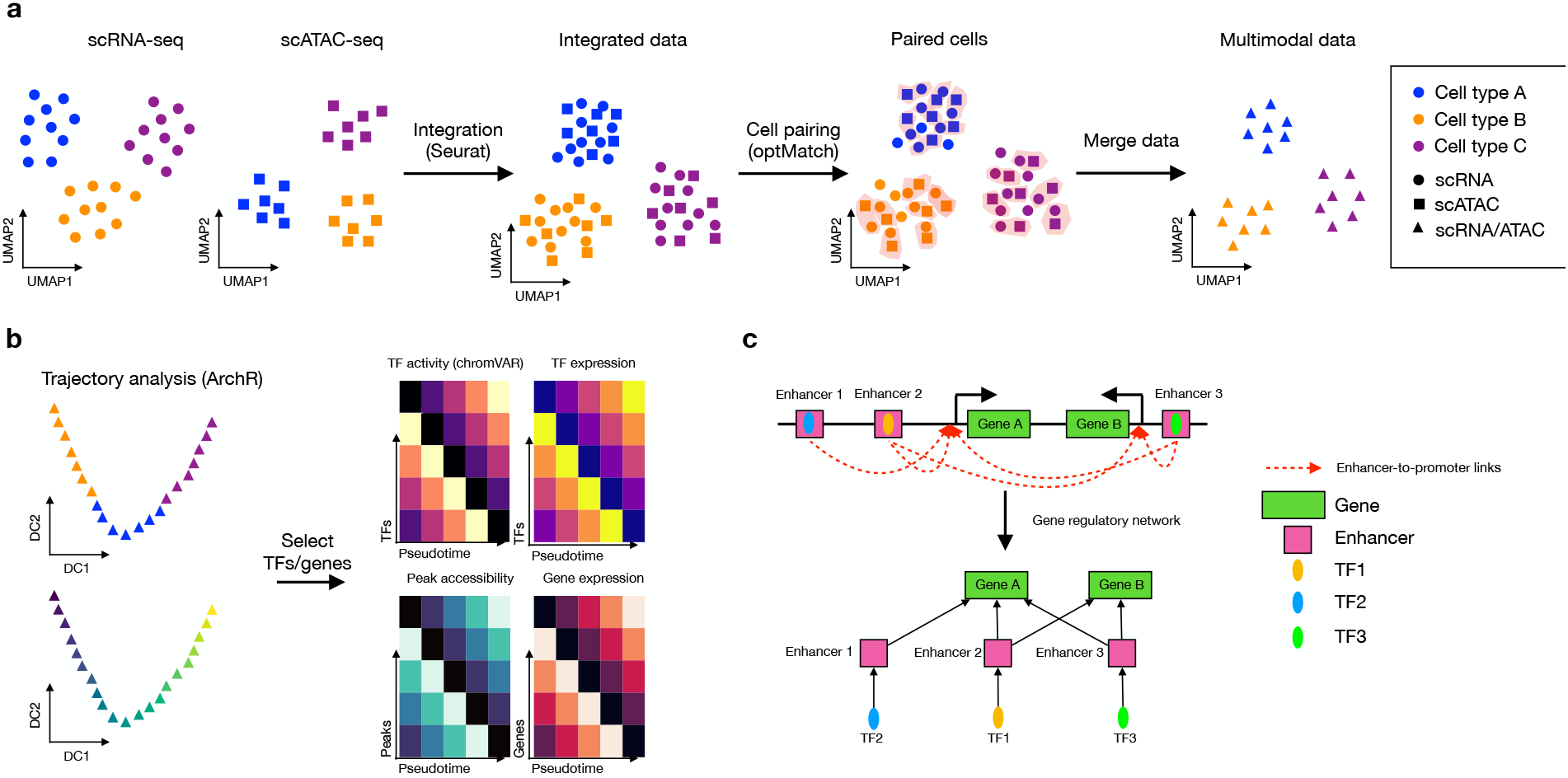
Overview of scMEGA. **a**, First step of scMEGA is to integrate the single-cell multi-omics profiles to obtain a paired dataset through modality integration built on the R package Seurat and cell pairing build on the approach OptMatch. The colors refer to different cell types and the shapes refer to different modalities. **b**, Next, scMEGA infers a pseudo-time trajectory to characterize the underlying dynamic process using ArchR. TFs are selected based on correlation analysis between binding activity (chromVAR) and expression along the trajectory. Genes are selected based on correlation analysis between peak accessibility and gene expression along the trajectory. **c**, Finally, scMEGA associates the selected TFs and genes to build a gene regulatory network based on predicted enhancer-to-promoter links.

## Results

### scMEGA identifies important regulators for myofibroblasts differentiation

We here provide a case study of using scMEGA to infer a GRN to understand the regulatory mechanism of fibrogenesis in human hearts after myocardial infarction [4]. We integrated the snRNA-seq and snATAC-seq data and identified four sub-populations of fibroblasts (Supplementary Fig. 1a). Detection of marker genes indicated that the cluster 2 highly expressed *SCARA5* which was recently reported as a marker for myofibroblast progenitors in the human kidney [11] (Supplementary Fig. 1b). The cluster 1 was marked by *POSTN, COL1A1*, and *COL3A1*, suggesting that these cells are differentiated myofibroblasts. Based on these, we built a pseudotime trajectory from cluster 2 to cluster 1 to study the myofibroblasts differentiation process (Supplementary Fig. 1c).

Next, we selected 79 candidate TFs and 2,207 genes that were used as input for inferring GRN (Supplementary Fig. 1d-e). Correlation between binding activity of the candidate TFs and expression of the candidate genes revealed two major regulation modules with each one corresponding to a distinct fibroblast subcluster (Supplementary Fig. 1f). Visualization of the inferred network pinpointed regulators of myofibroblast differentiation (Fig. 2a). For example, we identified that NR3C2 as a regulator of the *SCARA5* + fibroblasts (module 1 - fibroblast progenitors). NR3C2 is an important target in the clinic to halt the progression of fibrosis in heart failure [12] and we observed an decreased binding activity, TF expression, and target gene expression of NR3C2 along the trajectory (Fig. 2b). Regarding the myofibroblasts, we detected several fibrosis relevant TFs as TEAD [13] and RUNX family genes (Fig. 2b). Of note, we have recently characterized the role of RUNX1 as playing an important role in kidney [2] and heart [4] fibrogenesis.

As an example of downstream features of the eGRNs, we extracted target genes (regulomes) for NR3C2 and RUNX1 and inspected the expression of the TFs and their regulomes in space (Fig. 2c). We could not detect clear expression patterns of the TFs in space due to sparsity and low expression values of these TFs in spatial transcriptomics (Fig. 2d). By exploring the regulomes (target genes) of NR3C2 and RUNX1, we observed that gradients and mutually exclusive spatial expression in defined cardiac regions of fibrotic responses, highlighting the power of scMEGA in delineating TF regulome expression in sparse spatial transcriptomics data.

**Figure 2.**
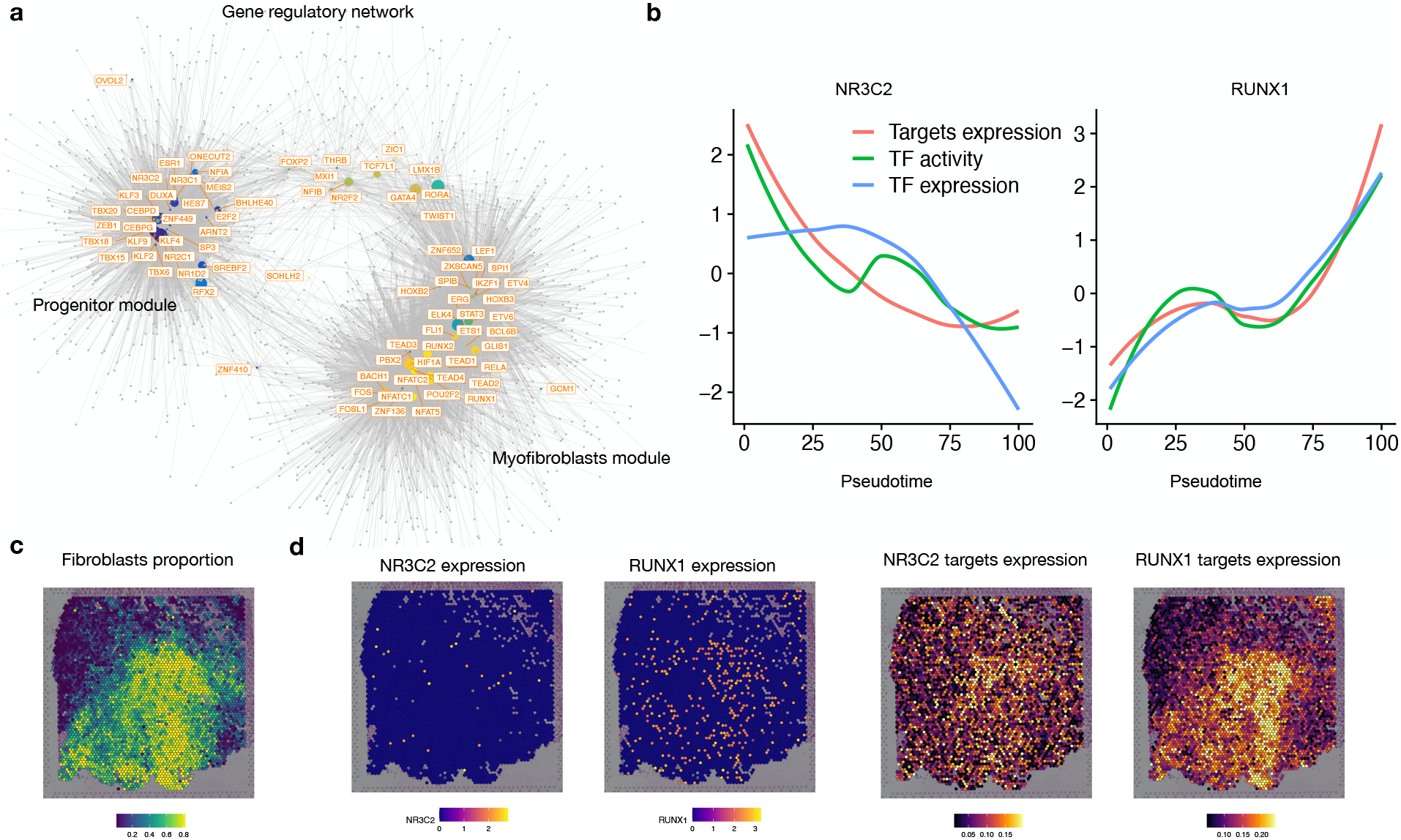
Case study of myofibroblast differentiation. **a**, Visualization of the inferred gene regulatory network for myofibroblasts differentiation. Each node represents a TF (regulator) or gene (target). TFs are colored by pseudotime point at which the TF has the highest activity score as estimated by chromVAR. **b**, Line plots showing the TF binding activity, TF expression and target expression along the myofibroblasts differential trajectory for NR3C2 and RUNX1. The x-axis represent the pseudotime points and the y-axis represent the z-score transformed values. **c**, Visualization of spatial distribution of fibroblast proportion estimated by cell2location in ischemic zone of human heart after myocardial infarction. **d**, Left: spatial distributed gene expression of NR3C2 and RUNX1. Right: spatial distributed gene expression of all targets of NR3C2 and RUNX1.

## Conclusions

In this work, we present scMEGA, a comprehensive and use-friend framework to infer enhancer-based GRN by using single-cell multi-omics profiles as input. scMEGA is built upon several popular R packages (e.g., Seurat, Signac and ArchR) for single-cell data analysis. It enables users to perform end-to-end GRN inferences and prioritize important TFs and genes for experimental validation and the use of regulomes to analyze spatial transcriptomics. We exemplify the use of scMEGA in fibroblasts in human hearts after myocardial infarction [4]. scMEGA is an important and unique framework for understanding complex gene regulation mechanism for various biological process from single-cell multi-omics data.

## Method

There are three major steps in scMEGA to infer a GRN, namely, (1) data integration, (2) identification of candidate TFs and genes, and (3) gene regulatory network assembly and visualization.

### Data integration

First step of scMEGA is to integrate the single-cell multi-omics profiles to create a pseudo-multimodal dataset where each cell is characterized by gene expression and chromatin accessibility (Fig. 1a). For integration, a gene score matrix estimated from scATAC-seq data is needed. This can be done by using the R package ArchR [7]. Next, scMEGA first uses the canonical correlation analysis (CCA) implemented by the R package Seurat [6] to project the cells into a shared low-dimensional space based on the gene expression from scRNA-seq and gene activity scores from scATAC-seq data. In case batch effects are present, the method Harmony [14] can be applied for batch correction.

Next, scMEGA performs cell pairing to obtain one-to-one matching between scRNA-seq and scATAC-seq, thus creating a pseudo-multimodal dataset that is used for downstream analysis. This step is built upon the OptMatch pairing approach proposed by Kartha et al. [15]. Finally, scMEGA merges the data for each of the pairs by extracting gene expression from scRNA-seq and chromatin accessibility from scATAC-seq. Of note, this step can be skipped in case single-cell multi-modal protocols (e.g., SHARE-seq [16] and 10X Multiome) were employed and a joint embedding of the modalities is available, such as provided by MOJI-TOO [17].

### Identification of candidate TFs and genes

scMEGA next identifies candidate TFs and genes based on the paired dataset which is obtained either through computational integration as described above or generated by using single-cell multimodal protocols. To do so, a pseudotime trajectory characterizing the underlying dynamic process is inferred by using the supervised approach as implemented by the R package ArchR [7]. Then, scMEGA estimates binding activity for each TF and cell using the package chromVAR [8] based on the predicted binding sites and chromatin accessibility profiles. To identify functionally relevant TFs for a biological process, scMEGA calculates the correlation between TF binding activity and TF expression. A high correlation indicates that the TF is both highly expressed and of which the motif is more accessible than the average profiles [3]. Next, scMEGA selects the candidate TFs based on statistical test of the correlation which can be defined by the user.

To select relevant genes, scMEGA first computes the expression variation of each gene along the pseudotime trajectory and picks up the top 10% (this cutoff can be adjusted by the user) most variable genes. To further identify the genes that are regulated by cis-regulatory elements (i.e., enhancers), scMEGA next predicts the enhancer-to-promoter links by computing the correlation between the chromatin accessibility of enhancers and the expression of nearby genes. Genes that are significantly associated with at least one enhancer are kept as targets. These analyses are performed based on functions from the ArchR [7], which were adapted to operate on Seurat objects.

Gene regulatory network construction and visualization Next, scMEGA builds a quantitative gene regulatory network by aggregating the information from enhancer-to-promoter links, predicted TF binding sites, and correlation of TF activity and target gene expression. More specifically, scMEGA first measures the correlation between binding activity of all the selected TFs and expression of all the selected genes as described above, which generates a quantitative regulation network. To link a TF to its target gene, scMEGA utilizes the predicted enhancer-to-promoter links and TF binding sites, i.e., a gene is only considered as the target of a TF if this gene is associated with at least one enhancer and this TF is bound to one of the associated enhancers, which creates a binary regulation network. By combining the quantitative and binary network, scMEGA produces a final gene regulation network prediction. Of note, this network can be further filtered by using the estimated correlation.

Next, scMEGA creates a directed (from TF to gene) and weighted (as measured by the correlation) graph based on the R package igraph [9] and prioritizes important TFs by computing the betweenness score and page-rank index for each TF [18]. Finally, scMEGA allows visualization of the network by exploring layout algorithms provided by igraph [9]. Additionally, when the spatial transcriptome (ST) data (e.g., generated by using 10X Visium assay) are available, scMEGA also allows to visualize target gene expression for selected TFs to understand the regulation activity in spatial coordinates.

## Acknowledgements

We would like to thank Ricardo O. Ramirez Flores for providing the pre-processed snRNA-seq data for fibroblasts and spatial transcriptomics slide and Julio Saez-Rodriguez for discussions on TF regulomes.

## Author’s contributions

ZL and ICG conceived the study. ZL implemented the software with help from JSN. CK and RK interpreted the analysis results for myofibroblasts differentiation. ZL written the original draft and ICG reviewed and edited the draft. All authors read and approved the final manuscript.

## Availability of data and materials

scMEGA is available from https://github.com/CostaLab/scMEGA. Markdown notebooks for the analysis of myocardial infarction data are found https://costalab.github.io/scMEGA/. The data used in this article are available in the Zenodo

Archive: https://zenodo.org/record/6623588.

## Ethics approval and consent to participate

Not applicable.

